# Membrane-MEDYAN: Simulating Deformable Vesicles Containing Complex Cytoskeletal Networks

**DOI:** 10.1101/2021.03.19.436229

**Authors:** Haoran Ni, Garegin A. Papoian

## Abstract

The plasma membrane defines the shape of the cell and plays an indispensable role in bridging intra- and extra-cellular environments. Mechanochemical interactions between plasma membrane and cytoskeleton are vital for cell biomechanics and mechanosensing. A computational model that comprehensively captures the complex, cell-scale cytoskeleton-membrane dynamics is still lacking. In this work, we introduce a triangulated membrane model that accounts for membrane’s elastic properties, as well as for membrane-filament steric interactions. The corresponding force-field was incorporated into the active biological matter simulation platform, MEDYAN (“Mechanochemical Dynamics of Active Networks”). Simulations using the new model shed light on how actin filament bundling affects generation of tubular membrane protrusions. In particular, we used membrane-MEDYAN simulations to investigate protrusion initiation and dynamics while varying geometries of filament bundles, membrane rigidities and local G-Actin concentrations. We found that bundles’ protrusion propensities sensitively depend on the synergy between bundle thickness and inclination angle at which the bundle approaches the membrane. The new model paves the way for simulations of biological systems involving intricate membrane-cytoskeleton interactions, such as occurring at the leading edge and the cortex, eventually helping to uncover the fundamental principles underlying the active matter organization in the vicinity of the membrane.

## 1 Introduction

Cytoskeletal fibers, along with regulatory molecules, such as α-actinin cross-linking nearby actin filaments, and myosin walking on the actin filaments, work together to establish cell’s main mechanochemical engine, continuously consuming energy. The plasma membrane, on the other hand, is mostly inactive. However, it defines the boundary of a cell, playing an important role in being a mechanical and chemical window to the outside world, regulating cytoskeletal functions [1].

Mechanically, the membrane is a thin fluid film, resisting in-plane and out-of-plane deformations, which, in turn, influences its cellular functions. For example, membrane tension has been shown to influence exocytosis, endocytosis and cell motility [2, 3]. Membranes can interact with the cytoskeleton mechanically via repulsion and adhesion [4]. Cells with the plasma membrane attached to a dense cross-linked actin cortex have their boundaries significantly rigidified [2]. On the contrary, disruption of filament-membrane attachments leads to cellular blebbing [5, 6]. Membrane proteins also participate in numerous signaling processes, regulating, in particular, many cytoskeletal interactions. For instance, the membrane attached WASp-VASP complex regulates Arp2/3 activation, which is an actin branching protein essential in lamellipodia formation [7].

Computer simulations can shed light on mechanisms of membrane-cytoskeleton interactions, building on and complementing extensive experimental observations of these processes [1]. A variety of such computational models have been proposed during the last two decades. Both analytical and computational models were used to study the filopodia formation and properties [8, 9]. To study lamellipodial protrusions, the membrane leading edge was modeled as a quasi 1D curve, interacting mechanochemically with a branched actin filament network [10]. Following up on these advances, it is desirable to develop a flexible, scalable, 3D model for membranes and their interactions with cytoskeleton that has detailed mechanochemical components essential for simulating whole-cell dynamics. With this goal in mind, we based our work on MEDYAN (Mechanochemical Dynamics of Active Networks, a software designed for carrying out detailed cytoskeletal simulations [11]), extending it such that cytoskeletal networks can be enclosed within a movable membrane domain.

MEDYAN can simulate cytoskeletal behaviors at high spatial and temporal resolutions. It couples cytoskeletal network mechanics with spatially resolved chemical dynamics, enabling studies of cytoskeletal processes at the micrometer scale and the time scale of tens of minutes [11]. Simulations using MEDYAN have shed light on the turnover and treadmilling dynamics for actin filaments [12], the stability and shape transformations of actin bundles [13] and the entropy production and cytoskeletal avalanches (cytoquakes) of evolving actomyosin networks that are far from equilibrium [14, 15, 16]. Currently, however, the enclosing domain for cytoskeletal evolution in MEDYAN is rigid and immovable, imposing inflexible steric restriction on cytoskeletal filaments. In this work, we incorporate into MEDYAN an explicit membrane model, which accounts for membrane’s salient mechanical properties, volumetric effects such as osmotic pressure, as well as filament-membrane interactions. The new membrane model paves the way for whole-cell simulations using MEDYAN.

In this work, we applied membrane-MEDYAN to simulate cytoskeletal networks enclosed in a vesicle, finding that actin network’s architecture plays an important role in generating membrane tubular protrusions. Furthermore, our analysis indicates that bundle protrusion initiation sensitively depends on local G-Actin concentration, bundle thickness, membrane rigidity and the inclination angle of the bundle approaching the membrane.

The remainder of the paper has the following organization. In Section 2, we introduce the new membrane model, including its discrete representation, the mechanical energy model and the membrane-cytoskeleton interactions. In Section 3, we discuss the application of membrane-MEDYAN to several relatively simple model systems. In particular, we report on our simulations of vesicles containing various actomyosin networks, exploring conditions that lead to formation of successful protrusions. Finally, in Section 4, we summarize our work, and further elaborate on the limitations and potential applications of the new model.

## 2 Methods

### 2.1 MEDYAN Simulations

The new membrane model was incorporated into MEDYAN. The latter can be used to simulate active matter systems using a compartment-based reaction-diffusion scheme, where diffusing chemical species are considered “well-mixed” in each compartment, but the locations for force bearing species (such as actin filaments and bound myosin motors) are explicitly modeled and tracked [11]. MEDYAN simulations are based on iteratively interleaving stochastic draws of chemical events, using the Gillespie algorithm, followed by a mechanical equilibration step. Importantly, chemistry influences mechanics and vice versa, creating intricate mechanochemical feedbacks.

### 2.2 Membrane mechanics and discretization

Typically, the phospholipid bilayer has a thickness of 3 - 7 nm [17]. At the scale of a whole cell (1 - 10 μm), the thickness of the plasma membrane is much smaller than its lateral dimensions. Therefore, we model the plasma membrane as a 2D Riemannian manifold, 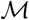, embedded in the 3D Euclidean space. Some of the mechanical properties of the lipid bilayer, arising from in-plane and out-of-plane deformations and from interactions with the cytoskeleton, are mapped onto this 2D manifold. We note here that from the perspective of a point on the surface, we use the term “lateral directions” to denote the directions in its tangent plane.

Next, we discretize the 2D surface 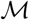 into a triangulated meshwork 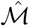 (Fig. 1a). The mesh is comprised from sets of vertices *V*, edges *E* and triangles *T*. Each vertex *υ*_*i*_ ∈ *V* is a representative point on the original surface. The 3D coordinates of all vertices control the shape of the surface. Each triangle *t* ∈ *T* is a set of 3 vertices {*υ*_*i*_, *υ*_*j*_, *υ*_*k*_} ⊂ *V*. These piecewise linear triangles tile the entire surface of 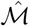, which, in turn, approximates the original manifold 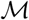. Each edge *e* is a set of 2 vertices {*υ*_*i*_, *υ*_*j*_} ⊂ *V*, and the set of all edges *E* ≔ {*e* = {*υ*_*i*_, *υ*_*j*_} : *e* ⊂ *t* for some *t* ∈ *T*}. We define the 1-ring neighbor vertices *N*_*υ*,1_(*υ*) ≔ {*υ*_*i*_ ∈ *V* : {*υ*, *υ*_*i*_} ∈ *E*}, and the 1-ring neighbor triangles *N*_*t*,1_(*υ*) ≔ {*t* ∈ *T* : *υ* ∈ *t*}. Several other constraints are added to the structure of the mesh to enforce a manifold-like structure. See the SI for more information. Below, all membrane related potentials take a discretized form as well, using coordinates of the vertices as corresponding arguments for the potentials and force functions elaborated below.

**Figure 1:**
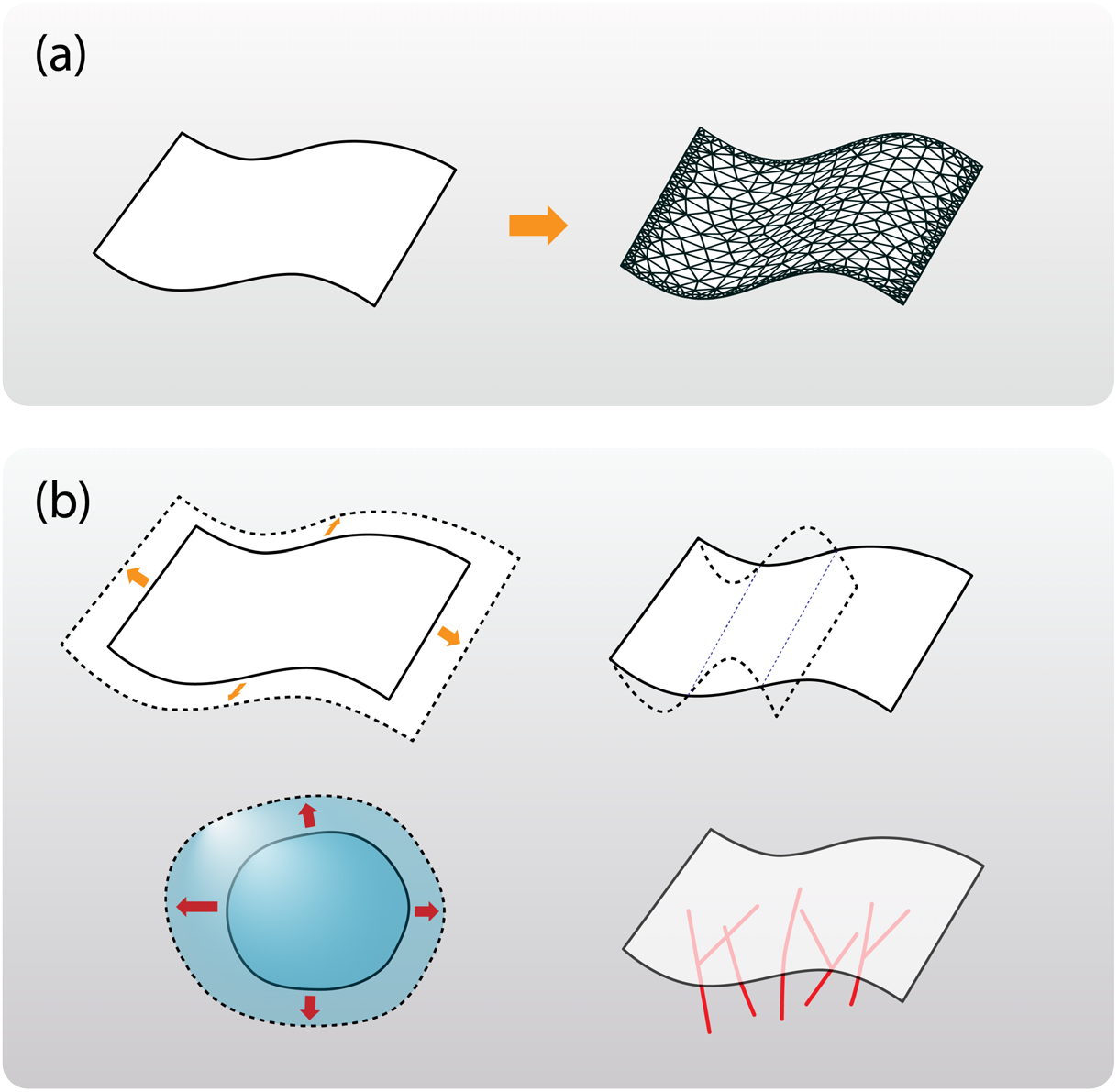
(a) The plasma membrane is represented as a 2D manifold 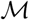 and is implemented as a triangulated mesh 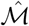. (b) The potential terms in our membrane model are illustrated. The shapes with a dashed stroke indicate allowed deformations that lead to: (top-left) surface tension; (top-right) membrane bending rigidity; (bottom-left) volumetric potentials in case of closed surface; (bottom-right) membrane-filament repulsion. The actin filaments are shown in red.

Excluding interactions with the cytoskeleton, we define the total mechanical energy for the membrane as the sum of the following potentials (see Fig. 1b top-left, top-right, bottom-left),

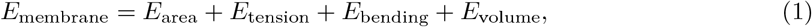

where *E*_area_ is the area elastic energy, *E*_tension_ is the surface tension energy, *E*_bending_ is the bending energy, and *E*_volume_ is the free energy associated with the 3D volume enclosed by the membrane, which can arise, for example, as a result of the intracellular osmotic pressure. The discretized form of these free energies are functions of mesh vertex coordinates, hence, one can compute the forces on vertices, with *f*_*ij*_ = −*∂E/∂x*^*j*^(*υ*_*i*_), where *f*_*ij*_ indicates the *j*-th component of the force on vertex *υ*_*i*_, *x*^*j*^(*υ*_*i*_) is the *j*-th coordinate of vertex *υ*_*i*_, for *υ*_*i*_ ∈ *V* and *j* ∈ {1, 2, 3}.

In the above outlined discretization scheme, mesh vertices do not have to represent the material points: the time evolution of an individual vertex does not represent the trajectory of any lipid molecule in the membrane. Therefore, one may also freely remesh the original surface at any time as long as the resulting shape is a good approximation of the original shape. Alternatively, we can enforce the vertices to be material points, where each vertex represents a patch of lipid molecules on the membrane that always stay together. In this case, the local area elasticity can be incorporated naturally in the membrane model, which will be discussed below. Our membrane model supports both options.

In addition to closed surfaces, we can also represent membranes with fixed boundaries. The boundaries can optionally connect the membrane surface to an implicit lipid reservoir, allowing the total number of lipid molecules to change.

#### Membrane area elasticity

A piece of free open membrane spontaneously relaxes to an equilibrium area that has zero surface tension [18]. Assuming homogeneity of lipid composition of the membrane, in the continuum limit, the membrane is considered to have a constant equilibrium mass density (mass per unit surface area) *ρ*_0_, so the free energy density (free energy per unit surface area) *e*_area_ can be written in a quadratic form

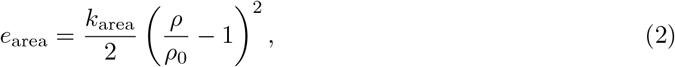

where *ρ* is the local density on the surface, and *k*_area_ is a positive constant with dimension MT^−2^, where M and T are the dimensions for mass and time, respectively. Total elastic energy is then, 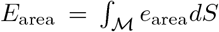. For a membrane not connected to a lipid reservoir, the mass conservation equation 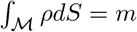 must be satisfied, where *m* is the constant total mass. In these integrals, *dS* is the surface area element.

Assuming that vertices represent material points, there is a natural discretization of surface elastic energy using Eq. 2. Each vertex *υ*_*i*_ carries a fixed amount of lipid molecules around it, and, therefore, has a well-defined equilibrium area *A*_0_(*υ*_*i*_) around it. The actual area around a vertex *A*(*υ*_*i*_) is defined as a third of the sum of areas of its 1-ring neighbor triangles, so 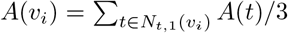, where *A*(*t*) is the area of triangle *t*. Assuming that the surface area density in the vicinity of a vertex is uniform, the area elastic energy of the discretized membrane can be written as

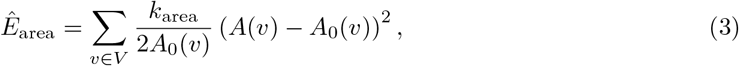

where the “hat” on *E* indicates that it is a discretized form.

On the other hand, if the vertices only parametrically represent the surface, not being connected to material points, the area elasticity cannot be naturally discretized around each vertex. However, if the membrane is not buffered by a lipid reservoir, we can use the mass conservation equation, with the assumption that the surface density is uniform across the whole membrane, to obtain the following relation,

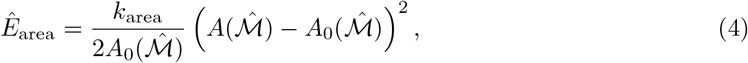

where 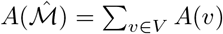 is the area of the discretized membrane, and 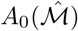 is the equilibrium area of the membrane. One can check, using the Cauchy-Schwarz inequality, that the energy in Eq. 4 is a minimum of the energy reachable in Eq. 3, if 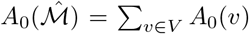. The uniform density assumption is valid, because at the smallest time resolution that we are interested in (~1 ms), the inhomogeneity of density on the membrane is quickly relaxed throughout the membrane due to its in-plane fluidity [2, 19].

In fact, the value of *k*_area_ for biomembranes is so large, that often the membrane area incompressibility is assumed, and the Lagrange multipliers are used in calculations to fix the total area [20]. However, since we employ energy minimization as the main mechanical simulation method, we use Eq. 3 or Eq. 4 to enforce a softer constraint, by allowing the area of membrane to change but with large energy penalty, instead of strictly fixing the area.

#### Membrane surface tension

Many types of cells can actively maintain the membrane tension within a certain range via various biological processes, such as membrane undulation and caveolae formation and dissolution, which function as implicit lipid reservoirs [2]. The experimentally measured tensions typically range from 0.01 mN m^−1^ to 0.04 mN m^−1^ [21].

If an implicit lipid reservoir is considered, the total mass of the membrane is no longer conserved, and in this case, the vertices in our model will not be associated with material points. Assuming a constant surface mass density and a constant chemical potential of the lipid reservoir, the total area-dependent energy is reduced to a linear form,

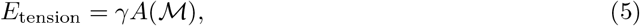

being further discretized to 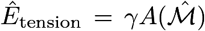 where *γ* is the surface tension, a property of the external reservoir.

#### Membrane bending elasticity

The bending energy can be described using the Helfrich Hamiltonian [22, 23],

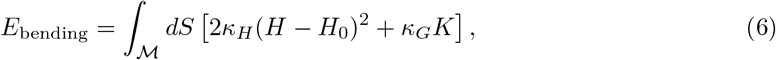

where *κ*_*H*_ and *κ*_*G*_ are the bending modulus and the saddle-splay modulus respectively, *H* is the mean curvature, *H*_0_ is the spontaneous curvature and *K* is the Gaussian curvature.

According to the Gauss-Bonnet theorem, the contribution from the second term (the Gaussian curvature term) is constant with fixed membrane topology and boundary conditions, so we can ignore the second term for most purposes. We discretize the first term in Eq. 6 on a per-vertex basis.

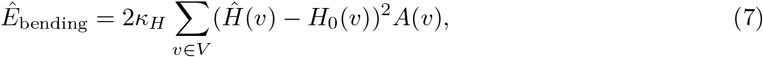

where 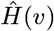 and *H*_0_(*υ*) are the mean curvature and the spontaneous mean curvature estimated around vertex *υ*, and are assumed to be uniform in the neighborhood of the vertex with area *A*(*υ*).

Many different ways of estimating the surface curvature on a triangulated surface mesh are discussed in [24, 25]. Based on our numerical experiments, we decided to allow for two different ways of estimating the mean local curvature at a vertex in MEDYAN, which provide good numerical stability for the systems that we studied. The first way is described in the Surface Evolver [26, 27],

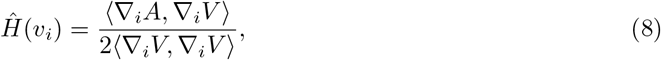

where *A* is the total area of the membrane, *V* is the total volume enclosed by the membrane, ∇_*i*_ means “taking the gradient with respect to coordinate of vertex *υ*_*i*_”, and (·, ·) denotes taking the inner product of the vectors. This definition gives a continuous and signed curvature value, the sign depending on the orientation of the surface, because the volume calculation is orientation-dependent. Therefore, this curvature estimate is suitable for bending energy computation when the spontaneous curvature *H*_0_ is not zero. In the continuum limit, ∇*A* and ∇*V* are both parallel to the normal direction, but computed numerically on a discretized mesh. Hence, there can be misalignments between these vectors. Under extreme conditions these two vectors can even become almost perpendicular, leading to unphysical geometries.

When *H*_0_ = 0, the sign of the curvature no longer matters in Eq. 7, and we rely on a way to directly estimate the unsigned curvature as,

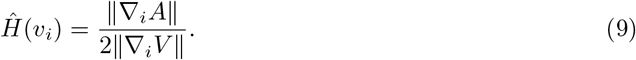

This equation provides better numerical stability under extreme conditions. For detailed discussion on how the bending energy was implemented, see the SI.

The volume for a closed discretized surface mesh can be computed as 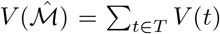, where *V* (*t*) is the signed volume of the tetrahedron formed by the triangle *t* and an arbitrary fixed point *p* ∈ ℝ^3^, and the sign depends on the orientation. It is often convenient to use the origin of the coordinate system as the fixed point.

#### In-plane shearing of the membrane is ignored

We do not explicitly include the in-plane shearing stress of the membrane in our model. Because the membrane behaves as a fluid in the lateral directions, the in-plane shearing stress cannot be sustained, while there can be a dynamic stress, which depends on fluid’s properties. In MEDYAN, mechanical energy minimization steps imply that the system is nearly equilibrated [11]. Therefore, we also ignore the dynamic shearing stress.

#### Volumetric energy

A membrane vesicle is also resistant to a change of its enclosing volume. The following quadratic expression captures the volume dependent free energy near the equilibrium volume,

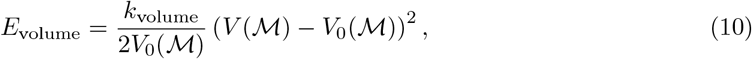

where 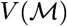 and 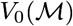 are the volume and the equilibrium volume enclosed by the membrane 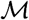 respectively, and *k*_volume_ is a positive constant, with dimension ML^−1^T^−2^, where M, L and T are the dimensions for mass, length and time, respectively. Next, we show how this formula can be derived using the osmotic pressure difference across the membrane vesicle.

Assuming a constant osmotic pressure outside the vesicle, the free energy due to osmotic pressure inside an enclosed volume is,

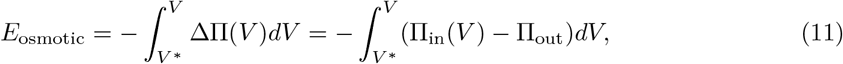

where Π_in_ and Π_out_ are the osmotic pressure inside and outside the vesicle respectively, ΔΠ = Π_in_ − Π_out_, *V* is the volume enclosed by the membrane and *V*\* is an arbitrary reference volume.

Under an assumption that the membrane is perfectly permeable to water and impermeable to solutes, the minimum osmotic energy occurs when *V* = *V*_0_ and Π_in_(*V*_0_) = Π_out_. Under further assumptions that the total mole fraction of the solutes is far less than the mole fraction of the solvent, and that the solutes are well mixed in the system, one can write Π_in_ = *Nk*_B_*T/V*, where *N* is the total number of solute molecules in the vesicle. In this case, *V*_0_ = *Nk*_B_*T/*Π_out_, and by choosing *V* * = *V*_0_, the energy can be evaluated as

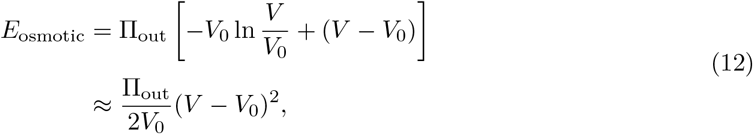

where the approximation is the result of expanding the energy to the second order with respect to *V* around *V*_0_.

#### Discussion of membrane force fields

For 2D manifold, all free energy terms mentioned above, except for the neglected in-plane shearing energy, are parametrization invariant, which means that the energy of the surface depends only on the shape of the surface. How the surface is triangulated or how the surface coordinates are chosen only affect the discretization error. More specifically, in the absence of membrane shearing energy, as long as the triangulations represent the same shape of the surface up to some precision (the similarity between two surfaces can be measured using the Hausdorff distance [28]), the total discretized free energies computed on those triangulations should be approximately the same. This is, in fact, very useful in a simulation, because it allows us to re-triangulate the membrane meshwork to improve the triangulation quality, without changing membrane’s physical behaviors. This approach effectively addresses the problem of the quality of triangulation becoming poor over time due to membrane’s large deformations, which, in turn, leads to undesirable numerical instabilities.

### 2.3 Adaptive surface remeshing

The aforementioned membrane potentials (especially bending energy, that is based on a discretized estimate for curvature) suffer numerical issues when the membrane deformation becomes large. To resolve this problem, we rely on the method of triangulated surface remeshing, which essentially optimizes the membrane quality without changing its shape [29, 30].

The quality of the mesh is related to the following: the triangle shape qualities and the dihedral qualities. For best numerical stability, the triangles used in the mesh should be as equilateral as possible, while the dihedral angles between the triangles sharing an edge should be as close to *π* as possible (or the angle between oriented normal vectors of the two triangles are as close to 0 as possible).

Our implementation of such an algorithm was inspired by [29, 30]. We apply local operations on the meshwork in an iterative manner to optimize both triangle and dihedral quality. With this algorithm, we are able to get an isotropic surface triangular mesh with good quality, where the vertex density adapts to the local curvature, making numerical simulations accurate for high curvature regions, while being efficient for low curvature regions. See the figures in the SI for an example of the remeshing procedure.

In MEDYAN, the adaptive mesh quality optimization for a membrane is carried out after every energy minimization. We rely on the Marching Tetrahedra algorithm as the mesh generator [31], which is fast, robust and easy to implement. However, the quality of the mesh by Marching Tetrahedra may lead to triangles that are far from being equilateral. Therefore, the mesh optimization algorithm is essential for achieving numerical stability.

### 2.4 Interactions between the membrane and the cytoskeleton

#### 2.4.1 Membrane-filament repulsion

Without tethering proteins connecting the cytoskeletal filaments to the plasma membrane, the filaments and the membrane sterically repel each other (Fig. 1b bottom-right). A hypothetical repulsive interaction between filament beads and membrane vertices would be insufficient in this case, because our simulations showed that the filaments are able to occasionally pierce through the triangle. Therefore, we developed an excluded volume interaction via repulsion between filament’s tip and a meshwork triangle, where essentially every point on the surface of the membrane interacts with the tip. The interaction energy between a given filament tip and membrane takes the following form,

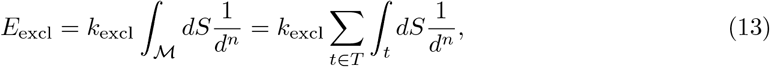

where *k*_excl_ is the force constant to the interaction and *n* describes the steepness of the potential, and *d* is the distance between the filament tip and a point on the membrane. This form of the potential generalizes the potential for segmented interactions from [32] to higher dimensional discrete elements. We found that choosing *n* = 4 produces a repulsive force field that is steep enough while still amenable to analytical evaluation of forces. For detailed derivation of this volume exclusion energy, see the SI.

The force acting on the filament tip from this repulsive interaction is *f* = −∇*E*_excl_, where ∇ means “taking the gradient with respect to the coordinate of the filament tip”. The reaction force (repulsive forces on the membrane by the filament tip) is balanced by the load on the membrane when the mechanical energy is minimized. Therefore, we directly use the value of *f* to be the load force when a filament polymerizes against a membrane. The polymerization of a filament near the membrane ratchets the membrane thermal fluctuation [33], and statistically work is done against the membrane load, resulting in membrane protrusion or deformation [34]. In the absence of explicit modeling of membrane fluctuation, we use the value of the load force *f* to compute a scaled filament polymerization rate [34, 11],

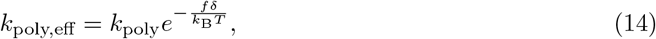

where *δ* is the effective monomer length (the average monomer length on the filament, and for actin, it is ~2.7 nm). This coupling introduces a non-linear feedback that allows instantaneously generated mechanical stresses to locally modulate chemical dynamics. A schematic diagram of membrane-filament interaction can be seen in Fig. 2.

**Figure 2:**
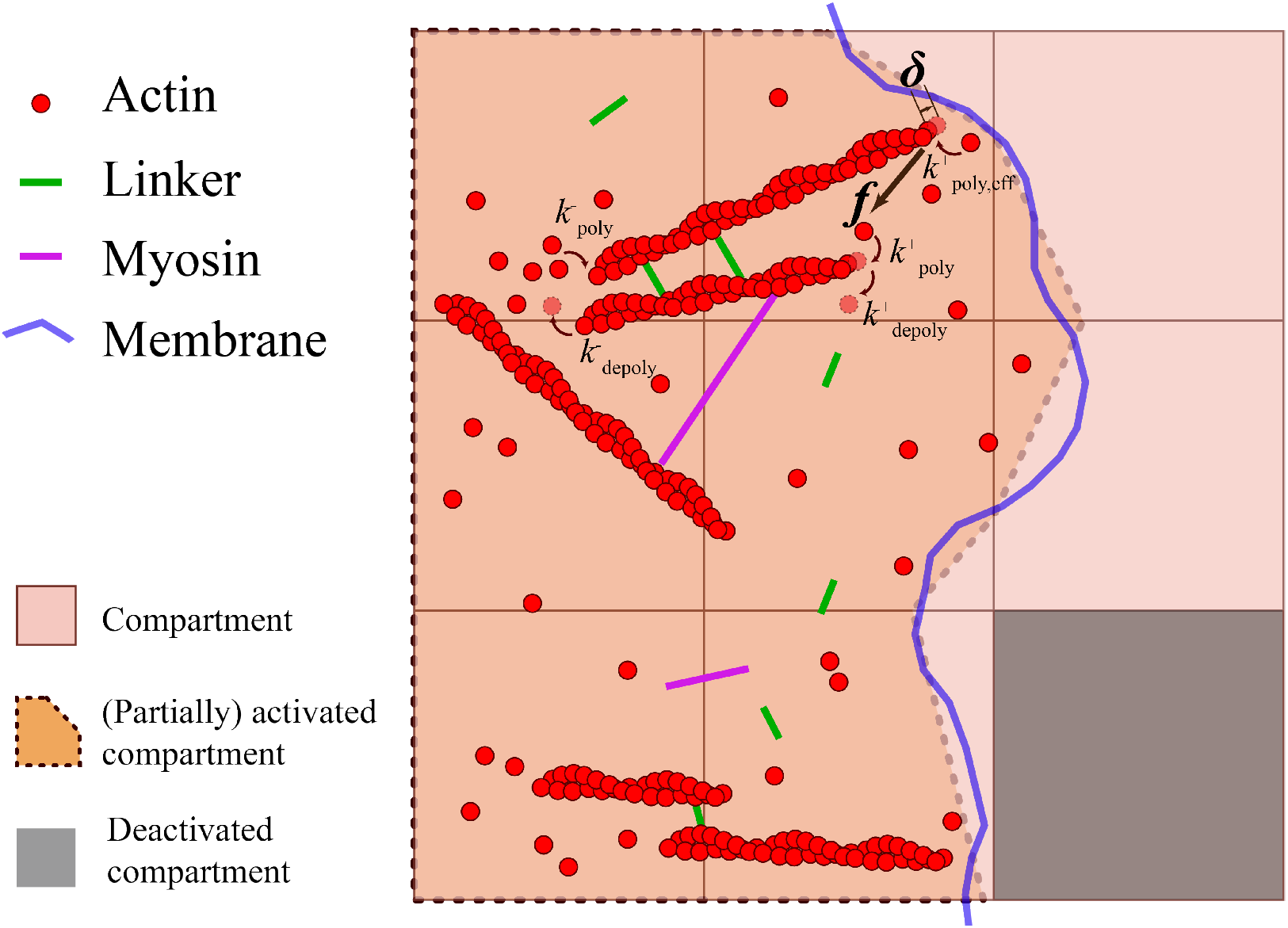
An illustration of membrane as a chemical regulator in 2D space. The membrane disallows big diffusing molecules to move across, such as G-Actin molecules or the α-actinin linkers. Therefore, the reaction volume is defined by the region inside the membrane (approximated by the straight dotted lines cutting through the compartments). The polymerization of filaments against the membrane is, however, possible but with reduced rates due to the Brownian ratchet effect. The actual implementation is carried out with 3D geometrical elements.

#### 2.4.2 Membrane and diffusing species

Diffusing large molecules, such as G-actin monomers, cannot move across the membrane without assistance. Therefore, the membrane must confine the space where the protein diffusion and chemical reactions take place. In a compartment based reaction-diffusion system, the effect of membrane confinement of reaction volume is achieved by full or partial deactivation of reaction networks in affected compartments. As shown in Fig. 2, a completely deactivated compartment does not allow chemical reactions or diffusion with neighboring compartments. A partially deactivated compartment, however, works as a normal compartment, but the reaction and diffusion rates are scaled by the diminished volume as well as the diminished contact area between neighboring compartments. To simplify computation, if a part of a membrane cuts through a certain compartment, we fit the shape of that part of the membrane into a plane, and perform a planar cut through that compartment to obtain the sliced volume and the sliced area on each side of the compartment.

## 3 Results and Discussion

We first applied the new force field to investigate how a spherical vesicle would deform as its volume starts to shrink under hyperosmotic conditions. The latter causes a mismatch of internal and external pressures, which affects our new volumetric term, leading to corresponding reduction of system’s volume. We started simulations with an initially spherical vesicle having a radius of 1 µm, applying potentials given in Eqs. 4, 7 and 12. The equilibrium area, *A*_0_, was fixed to be the initial spherical surface area. We continuously decreased the equilibrium volume *V*_0_ over time, which effectively creates an increasingly hyperosmotic chemical environment, since *V*_0_ = *Nk*_B_*T/*Π_out_ under dilute conditions.

Several snapshots of the simulation trajectory are shown in Fig. 3, indicating that under a constant area constraint, the membrane is unable to maintain its original spherical shape, crumpling as the vesicle volume shrinks. The local surface deformations, however, are smooth, because the bending potential prevents formation of sharp features where the mean curvature would be too large. We also notice that, over time, the regions with high convex curvature (as shown in blue in Fig. 3) become more distinct and highly localized.

**Figure 3:**
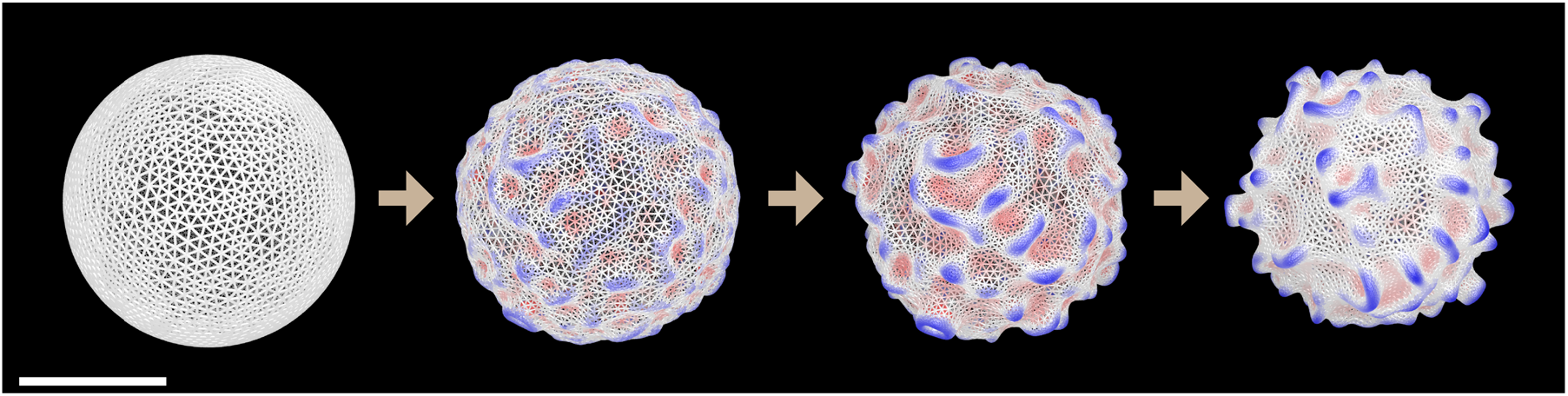
Evolution of the vesicle shape is shown under increasingly hyperosmotic conditions. Scale bar: 1 ìm. The red-white-blue coloring scale marks the local mean curvature, with red being the most concave and blue being the most convex. The membrane surface becomes more crumpled over time, and the bending stress is balanced by the forces arising from volume and area potentials. Using adaptive remeshing, the mesh is sampled more densely where the absolute values of local curvatures are higher.

Next we investigated how two classes of actomyosin architectures interact with the vesicle membrane boundary. Our simulations of vesicles containing disordered actomyosin networks, which are not bundled, indicate that randomly initialized filaments growing toward the membrane tend to bend instead of making a membrane protrusion, continuously inducing small membrane ruffles (Fig. 4a). At a steady state, the filaments accumulate just beneath the membrane surface. On the other hand, as displayed in Fig. 4b, the initially bundled filaments grow toward two poles of the vesicle, forming strong tubular protrusions. Interestingly, in some cases, a few filaments grow away from the axis of the bundle, become eventually buckled, hence, resembling the behavior of filaments in disordered actomyosin networks. Finger-like membrane protrusions, such as the filopodia, are ubiquitous structures in eukaryotic cells [35, 36]. Although the biological functions of the filopodia were widely explored [37, 36, 38], a clear understanding of how filopodia form as a result of membrane-cytoskeleton interactions is still lacking. Overall, these results demonstrate that distinct seeding patterns lead to different actomyosin network architectures, where formation of bundles is crucial for generating tubular membrane protrusions.

**Figure 4:**
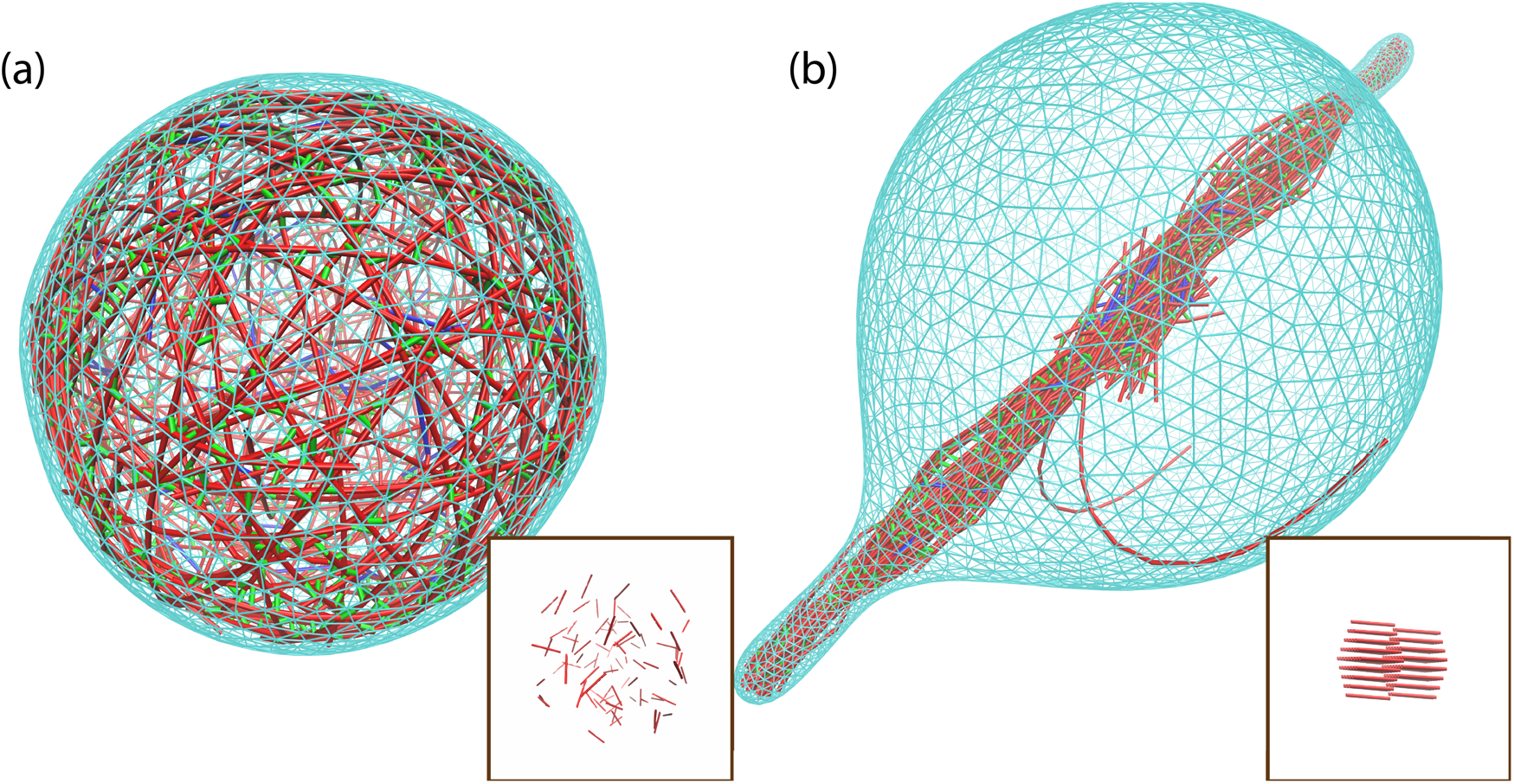
Vesicle deformation by actin filaments crucially depends on the spatial organization of actin filaments. The color codes are the same for two images: Cyan: Membrane, Red: Actin filaments, Green: α-actinin cross linker, Blue: Myosin motor. Filaments are randomly initialized in (a), and initialized as a bundle in (b). The boxed images on the bottom right show the initial filament arrangements for each condition. Bundled filaments can generate tubular protrusion, while filaments in disordered actomyosin networks accumulate under the membrane, being largely parallel to the surface.

To study what bundle properties most facilitate protrusion, we explored how the tubulation process depends on the number of actins filaments and G-Actin concentrations. To sustain stable tubular growth, the number of filaments, *N*, should satisfy 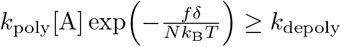 [8, 9], where 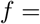 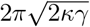, which is the load force at the tip of a stable filopodium. This imposes a minimum G-Actin concentration threshold which increases exponentially as the number of actin filaments decreases. For example, a bundle with 7 actin filaments requires at least 3.9 × 10^−4^ mol/m^3^ of G-Actin at the tip to sustain protrusion against the membrane with γ = 0.02 pN nm^−1^ and *κ* = 100 pN nm, but with 1 filament alone, the concentration has to be at least 0.474 mol/m^3^. Furthermore, for a tube to form in the first place, the filaments need to overcome the initial resistive force, which is greater in magnitude than subsequent steady state load forces. Another salient requirement for successful protrusion formation is for the bundle to avoid buckling [8].

To elaborate on these ideas, we simulated an actin bundle polymerization against a sheet of planar membrane at a constant surface tension with fixed edge boundaries. The results are summarized in Fig. 5. Our simulations show that a bundle with 3 filaments cannot effectively protrude against the membrane because it is not strong enough to resist buckling (Fig. 5a/b). With more actin filaments comprising the bundle, the protrusion can form at sufficiently high G-Actin concentrations (Fig. 5c/d). When a finger-like protrusion is successfully formed, its rate of growth increases with both the number of filaments in the bundle, and the G-Actin concentration (Fig. 5e).

**Figure 5:**
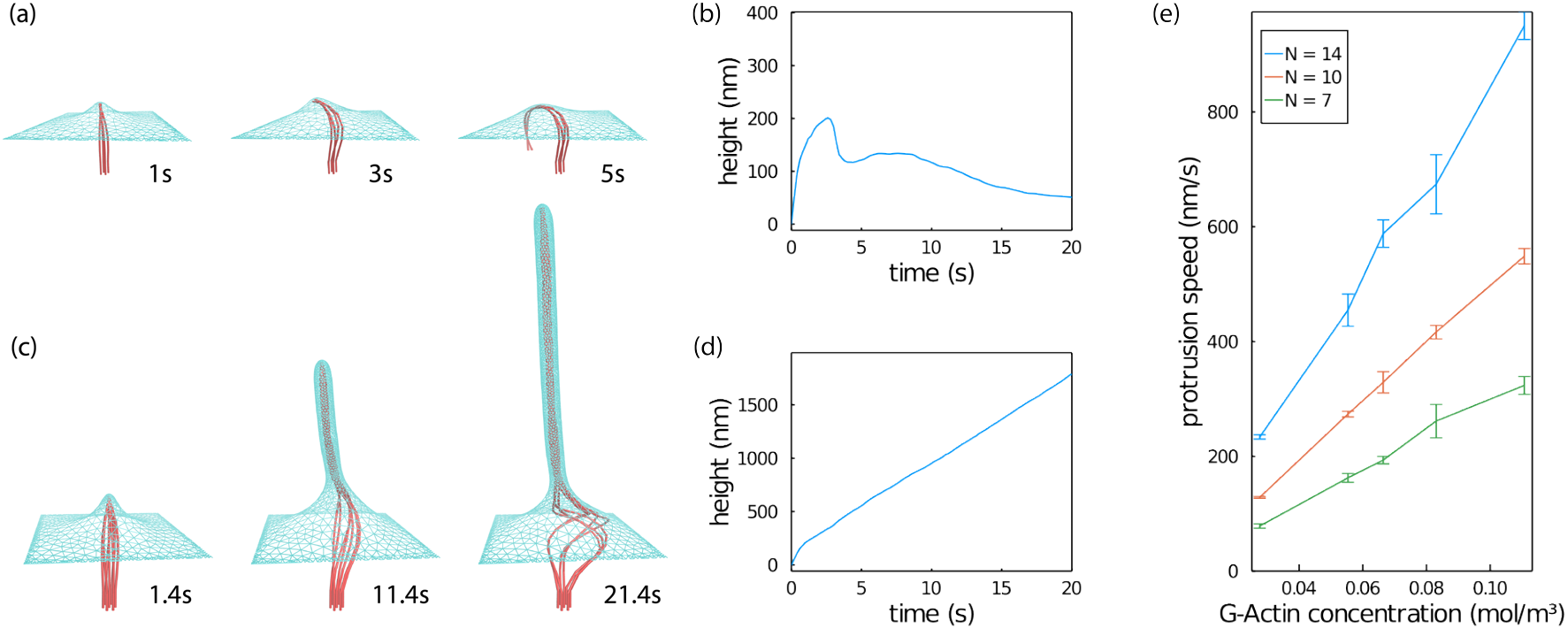
Filament bundle deforming the membrane. Color codes for (a) and (c): Cyan: Membrane, Red: Actin filaments. (a) An actin bundle with 3 filaments cannot make a successful protrusion. (b) The maximum height of the membrane corresponding to (a) is shown. The growing filaments slightly deform the membrane in the initiation phase, subsequently becoming buckled and, hence, failing to form a tubular protrusion. (c) A bundle with 7 filaments form a finger-like membrane protrusion. (d) The protrusion height on the membrane corresponding to (c) is shown. Interestingly, bundles tend to buckle less when surrounded by the membrane, in agreement with a previous work [43]. (e) The growth rate for a nascent tubular protrusion increases almost linearly as the G-Actin concentration increases, and also increases with the number of filaments in the bundle.

We assumed in this work that the pointed ends of the filaments are fixed, so there is no overall retrograde flow [39] of actin filaments. We also assumed that the G-Actin concentration is uniform along a membrane tube. Under these assumptions, the dependencies of the growth rate on the number of filaments in the bundle and the G-Actin concentration are qualitatively in accord with previous theoretical models [8, 40]. However, these assumptions allow the filopodia to grow at a constant speed indefinitely, which is unphysical in real filopodia. The existence of actin retrograde flow and the finite diffusion rate of G-Actin molecules lead to a steady state filopodial structure having a finite length and a dynamically generated G-Actin concentration gradient along the tube [9]. Furthermore, transportation of G-Actin by motor molecules, such as Myosin-X, may lead to a non-monotonic G-Actin concentration profile along the filopodium [41, 42]. These additional aspects of growth of mature filopodia are potentially amenable to simulations based on further elaboration of our new model.

In real cells, the polymerization directions of filament bundles are not necessarily perpendicular to the membrane. At sufficiently high G-Actin concentrations, we anticipate that the tubular protrusion is less likely to form when the filaments are inclined to the membrane surface, in particular, because those filaments would be more susceptible to buckling.

To investigate how bundle’s orientation affects protrusion formation, we simulated a filament bundle polymerizing towards a planar membrane sheet, where we varied the incident angle, *α*, which is the angle between the axis of the filament polymerization and the normal vector of the membrane. We also varied the load force at the tip of a stable protrusion, by modifying both the membrane tension and the bending rigidity simultaneously, while keeping their ratio constant, i.e. *γ* = χ*γ*_0_, *κ* = χ*κ*_0_ where *γ*_0_ and *κ*_0_ are the reference membrane tension and bending rigidity, and χ is a unitless parameter. In this way, the tubes formed would have the same radius 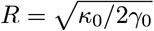 [8], but the load force at the tip for a stable protrusion changes as *f* = χ^2^*f*_0_. The results are demonstrated in Fig. 6. When the filament bundle is perpendicular to the membrane surface (*α* = 0), as the membrane rigidity increases (with increasing χ), the filaments first bend, and then eventually buckle, resulting in a protrusion failure. Furthermore, our simulations show that when filaments are not perpendicular to the membrane (*α* > 0), they easily buckle even under low membrane rigidities.

**Figure 6:**
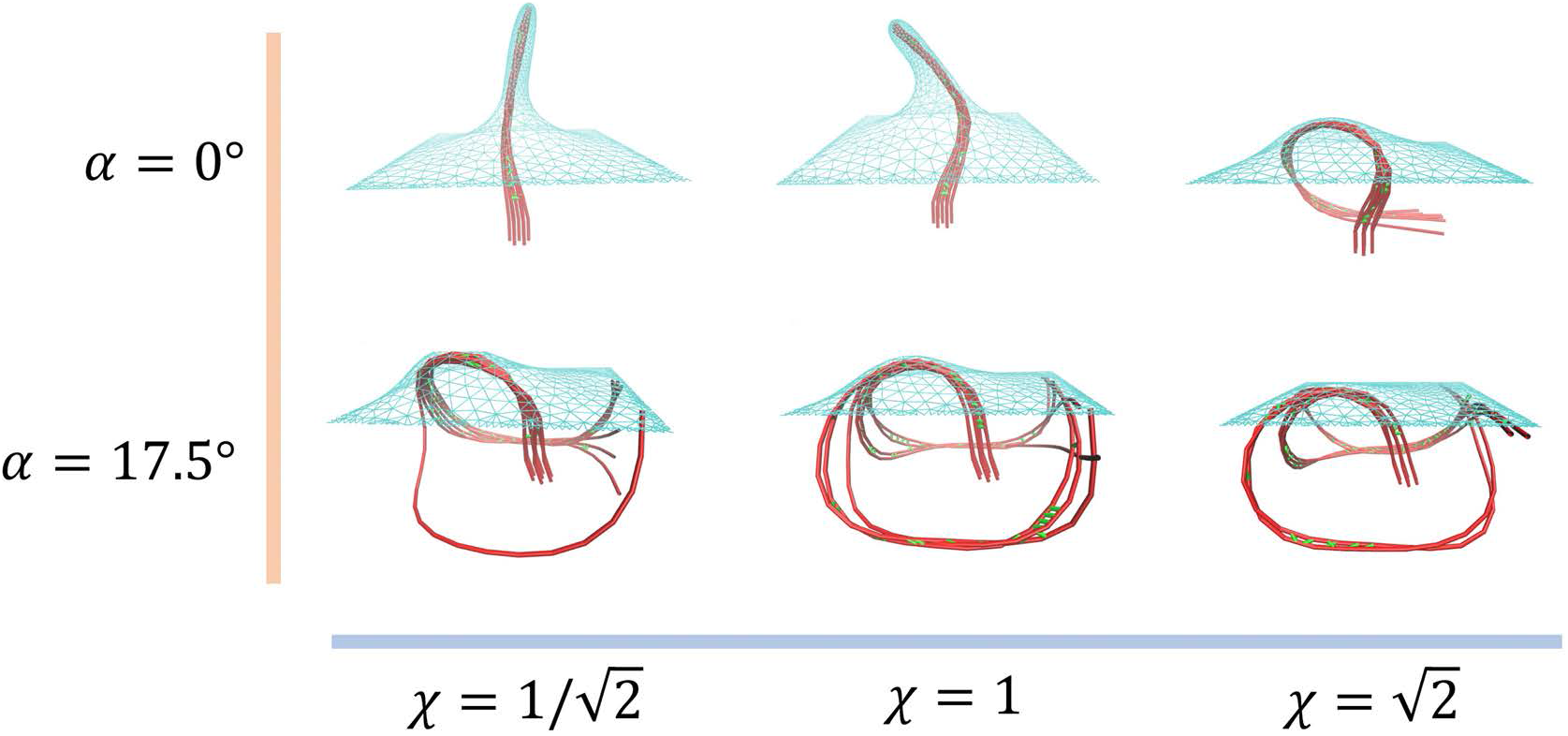
Membrane protrusion behaviors are shown under different relative membrane rigidities characterized by a unitless parameter χ (where a larger χ means larger membrane tension and bending modulus) and the incident angle α between the filament bundle and the membrane surface normal. All protrusions are made by a bundle comprised of 7 actin filaments. Color codes: Cyan: Membrane, Red: Actin filament, Green: α-actinin cross-linker.

The above-discussed results suggest that bundle’s orientation relative to the membrane is important in controlling the process of protrusion initiation. These insights help to explain the prior experimental observations showing that bundles growing within a vesicle can either be bent by the membrane and form ring-like structures, or protrude the membrane to form finger-like structures [44].

## 4 Conclusions

In this work we have addressed the problem of the interactions between the active matter and the plasma membrane, which are important for cell functions and have been a growing field of study. To study the rich behaviors arising from these mechanochemical interactions, we have developed a new force field for simulating the plasma membrane and its interaction with the active network. These algorithms were incorporated into MEYDAN, which is a broad simulation platform for active matter. The new membrane model enables simulations of rich cytoskeletal behaviors near the cell membrane, accounting for geometrical, mechanical and chemical details. On a single CPU, membrane-MEDYAN simulations can treat systems on the length scale up to several micrometers and time scales on the order of thousand seconds. Our software implementation allows to easily tune various model parameters, such as the elastic coefficients characterizing a membrane, and its initial shape. Our new model enables simulations of various cellular systems where membrane-cytoskeleton interactions play a salient role, such as lamellipodia, filopodia and dendritic spines.

In the current work, we applied membrane-MEDYAN to simulate a membrane vesicle, as well as the membrane-cytoskeleton systems under a variety of conditions. Simulations of an empty membrane vesicle under hyperosmotic stress lead to membranes’ intricate crumpling. Simulations of vesicles containing actomyosin networks show that bundling of actin filaments is crucial for generating tubular membrane protrusions. We found that the initial growth rate of successful protrusions is determined by the thickness of the protruding actin bundle and the G-Actin concentrations at the growth tip. Furthermore, unsurprisingly, lowering membrane’s surface tension or bending modulus facilitates the tubulation process. Finally we showed that the inclination angle between the filaments and the membrane is also critically important for the formation of finger-like protrusions. However, softer membranes can significantly mitigate this constraint.

One limitation of the current membrane model is the absence of membrane-associated proteins and membrane-cytoskeleton adhesions, which modulate both mechanical and chemical properties of the membrane-cytoskeleton system. Membrane based proteins, such as the BAR-domain proteins, can cause membrane remodeling by generating local membrane curvatures, being present in many cellular processes such as membrane trafficking and endocytosis [45, 46]. Membrane-cytoskeleton adhesions could increase the apparent rigidity of membrane attached to the actin cortex, which is important in cell blebbing [1, 4, 46]. They are also important in creating membrane invaginations, which are important in initiating endocytosis [47]. Moreover, chemical signaling via membrane receptors are known to regulate cytoskeleton behaviors [48]. Our current model can be naturally extended in future to enable studying of such intricate processes.

## Supporting information

Vesicle Containing Actin Bundle

Vesicle Containing Random Actin Filaments

## Acknowledgements

This work was supported by the grant CHE-1800418 from the National Science Foundation.

## Notes

### Competing Interest Statement

The authors have declared no competing interest.

