## Supplemental Information for "Membrane-MEDYAN: Simulating Deformable Vesicles Containing Complex Cytoskeletal Networks"

Haoran Ni and Garegin A. Papoian

March 15, 2021

#### 1 Simulation parameters and initial setups

Table 1 lists the default parameters used for simulation for this work. Unlisted parameters used in cytoskeleton simulation, unless otherwise stated, take the same value as in [1]. The membrane elastic parameters are chosen based on a prior study [2].

Table 1: The parameters used in simulation.

| Description | Symbol | Value |
|---|---|---|
| Membrane tension | $\gamma$ | $0.02\,\mathrm{pN\,nm^{-1}}$ |
| Membrane bending rigidity | $\kappa$ | $100\,\mathrm{pN\,nm}$ |
| Membrane spontaneous curvature | $H_0$ | 0 |
| Membrane area elasticity | $k_{\mathrm{area}}$ | $400\,\mathrm{pN\,nm^{-1}}$ |
| Membrane-filament bead repulsion constant | $k_{\mathrm{rep}}$ | $1300\,\mathrm{pNnm^3}$ |

##### 1.1 Simulations for interactions between actin and membrane vesicle

A simulation is set up for a vesicle system of radius $0.8\,\mu\mathrm{m}$ with the parameters in Table 2. The seeding actin filaments are 81 linear filaments with 80 monomers each, and they are initialized in two different ways. In the first case, the seeding filaments are initialized within the vesicle with random position and orientation. In the second case, the seeding filaments are initialized as a bundle at the center of the vesicle, with the barbed ends pointing at two poles of the vesicle (with 41 pointing to one pole and 40 pointing to the other pole), and with the pointed ends lying on a plane at the center. The membrane surface has constant surface tension, and the bending and osmotic potentials are included.

Table 2: Parameters for vesicle bundle deformation simulation.

| Description | Value |
|---|---|
| Actin monomer concentration | $0.1\,\mathrm{mol/m^3}$ |
| α-actinin cross-linker concentration | $5\times10^{-3}\,\mathrm{mol/m^3}$ |
| Myosin motor concentration | $5\times10^{-5}\,\mathrm{mol/m^3}$ |

##### 1.2 Perpendicular bundle protrusion against the membrane

The simulation starts with a planar sheet of membrane of size $1\,\mu\mathrm{m}\times1\,\mu\mathrm{m}$, with boundary vertices fixed and a constant surface tension of $0.02\,\mathrm{pN\,nm^{-1}}$. The bundled filaments are initially perpendicular to the surface of the membrane, each with length 432 nm, and the filaments are separated by a nearest distance of 40 nm. The barbed ends of the actin filaments point towards the membrane, and their the initial distances from the barbed ends to the membrane are 48 nm. We denote the direction of barbed end polymerization as the $+z$ direction. The segments at the pointed ends of the actin filaments are fixed in the space to provide support.

##### 1.3 Non-perpendicular bundle protrusion against the membrane

The simulations are initiated with a bundle of 7 filaments spaced 45 nm apart. The initial lengths of the filaments are 432 nm and the angle $\alpha$ between the direction of the filament and the normal vector of the membrane varies. The concentration of the G-Actin monomer is fixed, and the simulation parameters are listed in Table 3.

Table 3: Parameters for filament bundle polymerization with different contact angles with the membrane

| Description | Value |
|---|---|
| G-Actin monomer concentration | $0.1\,\mathrm{mol/m^3}$ |
| α-actinin cross-linker concentration | $2\times10^{-3}\,\mathrm{mol/m^3}$ |
| Reference membrane tension ($\gamma_0$) | $0.02\,\mathrm{pN\,nm^{-1}}$ |
| Reference membrane bending rigidity ($\kappa_0$) | $100\,\mathrm{pN\,nm}$ |

#### 2 Representation of membrane surface using a triangular mesh

We discretize the 2-dimensional surface $\mathcal{M}$ into a triangulated meshwork $\hat{\mathcal{M}}$. The elements in the mesh include sets of vertices $V$, edges $E$ and triangles $T$, which are described in the main text. The 1-ring neighbor vertices $N_{v,1}$ and the 1-ring neighbor triangles $N_{t,1}$ are also defined in the main text. Two triangles $t_i$, $t_j$ are called *adjacent* if $t_i \cap t_j \in E$. Now, given a vertex $v \in V$, if we define a graph with the nodes being elements in $N_{t,1}(v)$, and two nodes are connected if and only if two triangles share the vertex $v$ and are adjacent, then this graph can be divided into a set of connected components, and for each connected component, the collection of nodes (triangles) is called a *fan* around the vertex $v$.

To enforce a manifold-like structure, more rules need to be enforced on the elements of $\hat{\mathcal{M}}$: (1) Every vertex must exist in at least 1 triangle. This also implies that every vertex exists in at least 2 edges. (2) Every edge must show up in no more than 2 triangles. By definition, an edge shows up in at least 1 triangle. We call the edge that shows up in exactly 1 triangle the *border edge*. (3) For any $v \in V$, there is exactly one fan around $v$. Here are some intuitions behind the rules. Rule (1) is to make sure that there is no vertex that is not "connected to" another vertex. Rule (2) prevents more than 2 triangles sharing the same edge, which leads to a non-manifold structure. Rule (3) is to ensure that each vertex exists at exactly one location on the surface $\hat{\mathcal{M}}$. Without it, Multiple locations on $\hat{\mathcal{M}}$ can share one vertex. See Fig. 1.

If the original manifold $\mathcal{M}$ is orientable, $\hat{\mathcal{M}}$ can also be defined with an orientation. All the element definitions are the same with the previous definitions except for the following notable changes. A triangle $t \in T$, instead of being a set of 3 vertices, is now defined as an equivalence class identified by 3 distinct vertices $[v_i, v_j, v_k]$, with the equivalence being the cyclic permutation of index $i, j, k$. In other words, $[v_i, v_j, v_k] = [v_j, v_k, v_i] = [v_k, v_i, v_j]$. However, we still mandate that the set of vertices in each triangle is distinct. For example, although $[v_1, v_2, v_3] \neq [v_1, v_3, v_2]$, at most one of the two is allowed to show up in $T$. In other words, the triangle can still be viewed as a set of 3 vertices, but with the additional information of the order of the vertices. Then, an edge is defined similarly as before (treating each triangle as a set of 3 vertices), but now it is also useful to define a set of directed edges (or half edges) $H$, each element being an ordered pair of distinct vertices $(v_i, v_j)$, so $H = \{(v_i, v_j) \in V \times V : \{v_i, v_j\} \in E\}$. Apparently, for each edge $\{v_i, v_j\}$, there are 2 corresponding half edges $(v_i, v_j)$, $(v_j, v_i)$. Now the rules to enforce a manifold-like structure applies as well, but another rule needs to be added to account for the orientation. A half edge $h = (v_i, v_j)$ is *associated* with a triangle $t$ if $t = [v_i, v_j, v_k]$ for some $v_k$. Rule (4): For any edge that shows up in 2 triangles, each of the half edge corresponding to the edge must be associated with exactly one of the triangles. The intuition for this rule is that, if two oriented triangles are adjacent, the oriented normal vectors

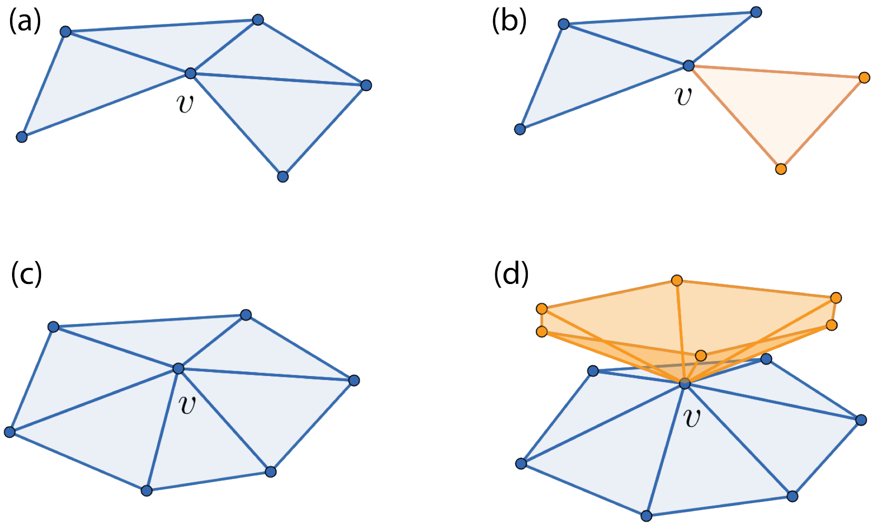


Figure 1: Examples of fans around the vertex $v$. Multiple fans are labeled using different colors. The configurations (a) and (c) each has one fan. The configurations (b) and (d) each has two fans, which is forbidden in the mesh.

poke out from the same side of the surface.

In this work, we use oriented triangular mesh to represent an orientable membrane surface. If a non-orientable surface is required, one can extend the existing orientable mesh to an oriented double cover of the non-orientable mesh, where the set of vertices in each triangle being distinct is no longer required.

### 3 Benchmark on membrane adaptive remeshing

See Fig. 2 for the result of a benchmark of our adaptive remeshing algorithm.

### 4 Membrane bending energy on discretized surface

The bending terms in the Helfrich Hamiltonian can be used for computing the bending energy of the membrane $\mathcal{M}$ [3, 4]

$$E_{\text{bending}} = \int_{\mathcal{M}} dS \left(2\kappa_H (H - H_0)^2 + \kappa_G K\right) \tag{1}$$

where $H$ is the mean curvature, $K$ is the Gaussian curvature, $H_0$ is the spontaneous curvature, $\kappa_H$ and $\kappa_G$ are the bending modulus and the saddle splay modulus respectively. Note that here the mean curvature $H$ stands for "the mean of two principal curvatures", while some other literatures use the term "mean curvature" as the local trace of the shape operator [5, 6], which is $2H$ in the context of this article.

For a membrane with fixed topology and boundary conditions, the contribution from the Gaussian curvature is a constant, so for most of our purposes we do not care about the second term in Eq. 1.

#### 4.1 Edge based curvature and bending energy estimation

When a smooth surface is represented by the discretized triangles, the curvature on the surface can be viewed as being concentrated on the edges [6]. Consequently, a natural way of computing the

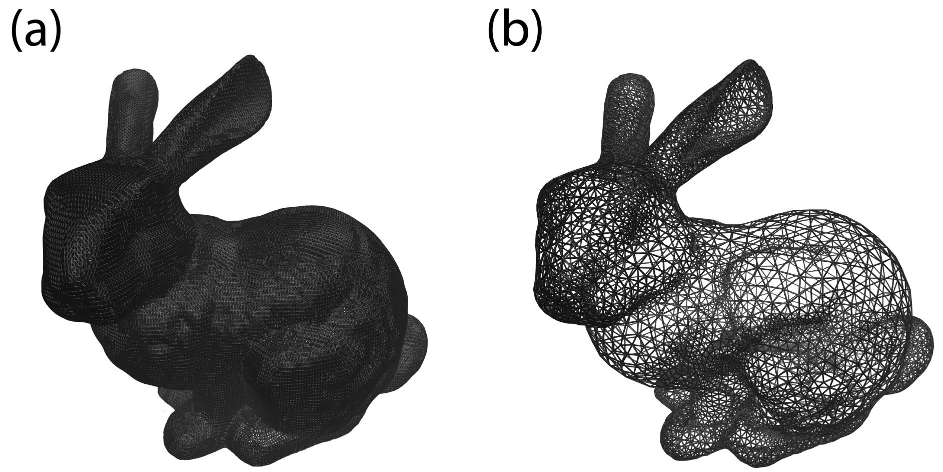


Figure 2: The remeshing algorithm generates anisotropic mesh with adaptable sizes suitable for simulation. Left: before remeshing; Right: after remeshing. (a) Showing the reconstructed mesh of the bunny model from the Stanford Computer Graphics Laboratory. The mesh is very dense and largely anisotropic. (b) The bunny mesh after the remeshing. The vertices are sparser than the original mesh, especially in relatively flat regions. Additionally, the vertex density increases in regions with higher local curvature.

discrete surface bending energy is to use the dihedral angle formula [7, 8, 9, 5].

$$\hat{E}_{\text{bending}} = \tilde{\kappa} \sum_{\alpha,\beta \in T} (1 - \langle \boldsymbol{n}_\alpha, \boldsymbol{n}_\beta \rangle) \tag{2}$$

where $\tilde{\kappa}$ is the force constant, $T$ is the set of all the triangles in the mesh, and $\alpha$ and $\beta$ are adjacent oriented triangles with unit normal $\boldsymbol{n}_\alpha$ and $\boldsymbol{n}_\beta$. It is one of the simplest ways to implement the bending energy, but the relation between the constant $\tilde{\kappa}$ and the real physical constants should be carefully considered.

It is shown [5] that under the continuum limit, Eq. 2 is equivalent to

$$E_{\text{bending}} = \kappa \int_{\mathcal{M}} dS \; (2H^2 - K) \tag{3}$$

which is the Helfrich energy when $H_0 = 0$, $\kappa_H = \kappa$ and $\kappa_G = -\kappa$. However, the relationship between $\kappa$ and $\tilde{\kappa}$ depends on the overall shape of the membrane [10], so for a membrane with arbitrary shape, this method might not converge to the Helfrich bending energy even at continuum limit. One of the origins of the difficulty is that while the discretized integral of mean curvature converges in the continuum limit, the discretized integral of squared mean curvature is singular [11, 10]. The integral of mean curvature is still useful, however, when the spontaneous curvature $H_0$ is not zero, and then a linear integral of mean curvature can be extracted from the Helfrich free energy and estimated based on dihedral angle between edges

$$\begin{aligned} E_{\text{bending,linear}} &= -4\kappa_H H_0 \int_{\mathcal{M}} dS \; H \\ &\approx -4\kappa_H H_0 \sum_{\alpha,\beta \in T} l_{\alpha\beta} \arccos(\langle \boldsymbol{n}_\alpha, \boldsymbol{n}_\beta \rangle) \end{aligned} \tag{4}$$

where $l_{\alpha\beta}$ is the length of the edge shared by neighboring triangles $\alpha$ and $\beta$, and the approximation comes from the discretization.

##### 4.2 Vertex based curvature and bending energy estimation

Despite the singularity of integral of squared mean curvature on a discretized triangular surface, one can still estimate the local mean curvature value around each vertex and use that to compute the bending energy [11].

$$E_{\text{bending}} = 2\kappa_H \int_{\mathcal{M}} dS\ (H - H_0)^2 \approx 2\kappa_H \sum_{v \in V} \hat{A}(v)(\hat{H}(v) - H_0(v))^2 \tag{5}$$

where $V$ is the set of all vertices, and $\hat{A}(v)$ and $\hat{H}(v)$ are the area and the mean curvature estimated around vertex $v$ respectively.

Many ways of estimating $\hat{A}(v)$ and $\hat{H}(v)$ exist [6]. Currently, in MEDYAN, the area around the vertex is a third of the sum of all the neighboring triangles sharing the vertex, and the curvature is estimated using the following formula [11, 12]

$$\hat{H}(v_i) = \frac{\langle \nabla_i A, \nabla_i V \rangle}{2\langle \nabla_i V, \nabla_i V \rangle} \tag{6}$$

where $A$ is the total area of the membrane, and $V$ is the total volume inside the membrane, which allows for continuous signed local curvature (signedness for curvature is important when $H_0 \neq 0$ in Eq. 5), and avoids numerical singularity when $\nabla_i A = 0$ on flat surfaces. In the continuum limit, $\nabla A \parallel \nabla V$ and are both along the normal direction, but numerically in a discretized mesh, there can be misalignments between the vectors mentioned above, and under extreme conditions these two vectors can even try to become perpendicular and produce unphysical geometries. A method with better numerical stability is by directly estimating the unsigned version of $\hat{H}(v_i)$

$$\hat{H}(v_i) = \frac{\|\nabla_i A\|}{2\|\nabla_i V\|} \tag{7}$$

This provides better numerical stability against several unphysical geometries in the first implementation. But in this case, we cannot directly incorporate the spontaneous curvature because the orientation information is lost in the equation. Nevertheless, the extra linear term can be extracted and computed in Eq. 4.

### 5 Derivation of triangle-bead volume exclusion interaction

The triangle-bead volume exclusion interaction takes the following form:

$$\hat{E}_{\text{excl}} = k_{\text{excl}} \int_{\hat{\mathcal{M}}} dS \frac{1}{d^n} = k_{\text{excl}} \sum_{t \in T} \int_t dS \frac{1}{d^n} \tag{8}$$

Here we derive the explicit analytical result when $n = 4$ and will discuss the geometric interpretation of some terms in the result.

##### 5.1 Evaluating the integral

The integration can be done as follows. Suppose we have a triangle $t$, with 3 vertices $v_0, v_1, v_2$ with Cartesian coordinates $x_0, x_1, x_2$. Let $r_{01} = x_1 - x_0$ and $r_{12} = x_2 - x_1$, any point $q$ on the triangle has the coordinate $x_q = x_0 + \alpha r_{01} + \alpha\beta r_{12}$, $\alpha \in [0, 1], \beta \in [0, 1]$, as displayed in Fig. 4. Using $\alpha$, $\beta$ as the surface coordinates, a metric tensor can be obtained naturally from the embedding in $\mathbb{R}^3$. In the matrix form, the metric tensor writes

$$(g_{\mu\nu}) = \begin{pmatrix} \|r_{01} + \beta r_{12}\|^2 & \langle \alpha r_{12}, r_{01} + \beta r_{12} \rangle \\ \langle \alpha r_{12}, r_{01} + \beta r_{12} \rangle & \|\alpha r_{12}\|^2 \end{pmatrix} \tag{9}$$

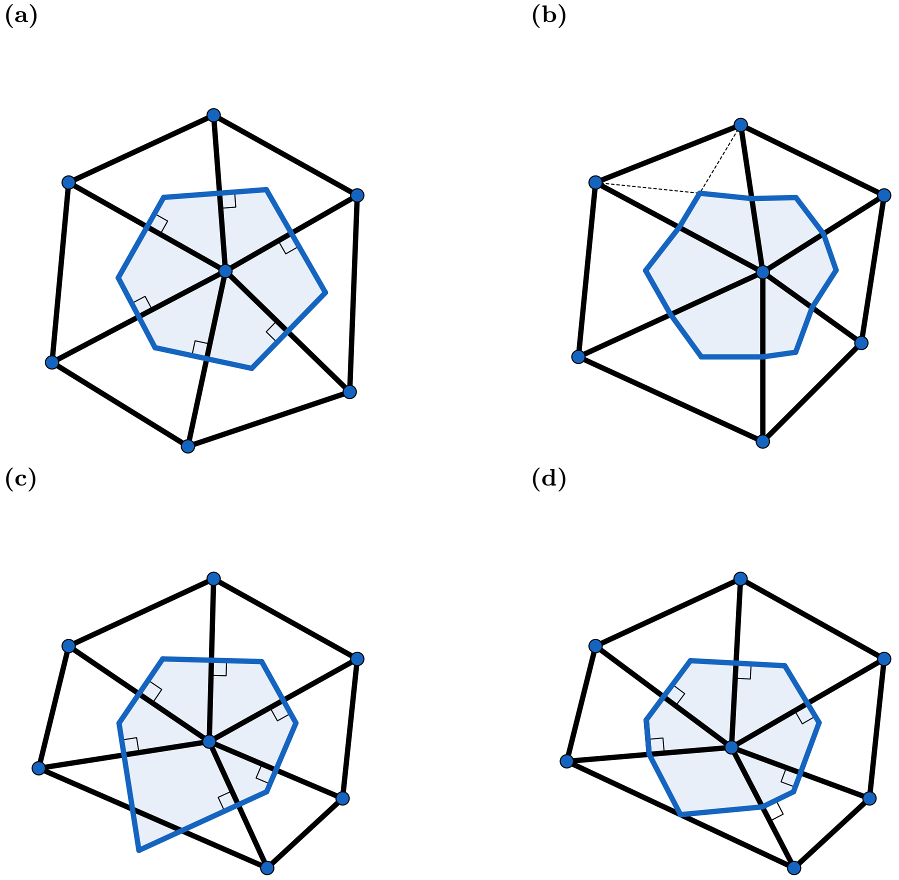


Figure 3: Ways of estimating the area at a vertex. The area is estimated from the region that is marked with blue shade. (a) Voronoi cell of the center vertex. (b) 1/3 of sum of area of all neighboring triangles around the center vertex. (c) Voronoi cell of the center vertex when obtuse triangles are involved. Therefore, area estimation using Voronoi cell only would suffer from great numerical errors when bad-quality triangles are involved. (d) Mixed area of the center vertex when obtuse triangles are involved [13].

where each element $g_{\mu\nu}$ is the inner product between tangent vectors $\partial_\mu$ and $\partial_\nu$, where $\mu, \nu$ can be $\alpha$ or $\beta$. Let $g$ be the determinant of the metric tensor. Then

$$\begin{aligned} g &= \|r_{01} + \beta r_{12}\|^2 \|\alpha r_{12}\|^2 - \langle \alpha r_{12}, r_{01} + \beta r_{12} \rangle^2 \\ &= (\alpha \|r_{01} \times r_{12}\|)^2 \end{aligned} \tag{10}$$

where $\|r_{01} \times r_{12}\|$ is simply twice the area of the triangle $A(t)$.

Suppose we have a filament tip $p$ at position $x_p$, and let $r_{p0} = x_0 - x_p$, then the distance between $p$ and $q$ is

$$d = \|\alpha r_{01} + \alpha\beta r_{12} + r_{p0}\| \tag{11}$$

The interaction energy can be written as

$$\hat{E}_{\text{excl}}(t, p) = k_{\text{excl}} \int_t dS \frac{1}{d^4} \tag{12}$$

where $k_{\text{excl}}$ is the strength of the interaction. The area element is the volume form on the surface, and can be expressed as

$$\begin{aligned} dS &= \sqrt{g}\; d\alpha \wedge d\beta \\ &= \alpha \|r_{01} \times r_{12}\| d\alpha \wedge d\beta \end{aligned} \tag{13}$$

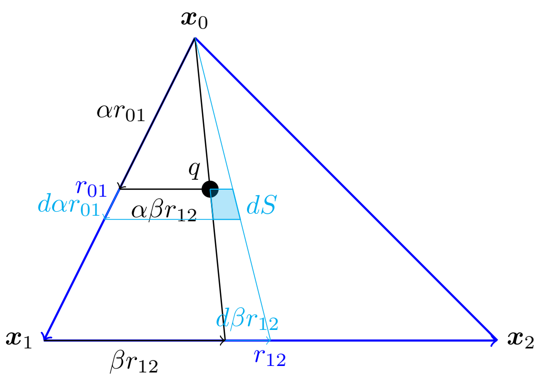


Figure 4: The area element on a triangle facet.

Now Eq. 12 can be written as

$$\hat{E}_{\text{excl}} = k_{\text{excl}} \|r_{01} \times r_{12}\| \underbrace{\int_0^1 \alpha\; d\alpha \int_0^1 d\beta\; \|\alpha r_{01} + \alpha\beta r_{12} + r_{p0}\|^{-4}}_{I} \tag{14}$$

Then by letting

$$\begin{aligned} a &= \|r_{01}\|^2 \\ b &= \|r_{12}\|^2 \\ c &= \|r_{p0}\|^2 \\ e &= \langle r_{01}, r_{p0} \rangle \\ f &= \langle r_{12}, r_{p0} \rangle \\ g &= \langle r_{01}, r_{12} \rangle \end{aligned} \tag{15}$$

one can get

$$I = \int_0^1 \alpha\; d\alpha \int_0^1 d\beta\; \left(\alpha^2 a^2 + \alpha^2 \beta^2 b^2 + c^2 + 2e + 2\alpha\beta f + 2\alpha^2 \beta g\right)^{-2} \tag{16}$$

we are able to evaluate the integral analytically with

$$
\begin{aligned}
I = \Bigg( & -\frac{(af-eg)\arctan\frac{e}{\sqrt{ac-e^2}}}{\sqrt{ac-e^2}} \\
& + \frac{(af-eg)\arctan\frac{a+e}{\sqrt{ac-e^2}}}{\sqrt{ac-e^2}} \\
& + \frac{(af+fg-be-eg)\arctan\frac{e+f}{\sqrt{ac-e^2-2ef-f^2+bc+2cg}}}{\sqrt{ac+bc-e^2-2ef-f^2+2cg}} \\
& + \frac{(be-af+(e-f)g)\arctan\frac{a+b+e+f+2g}{\sqrt{ac-e^2-2ef-f^2+bc+2cg}}}{\sqrt{ac+bc-e^2-2ef-f^2+2cg}} \\
& + \frac{(ab+be-g(f+g))\arctan\frac{f+g}{\sqrt{b(a+c)+2be-(f+g)^2}}}{\sqrt{b(a+c)+2be-(f+g)^2}} \\
& - \frac{(ab+be-g(f+g))\arctan\frac{b+f+g}{\sqrt{b(a+c)+2be-(f+g)^2}}}{\sqrt{b(a+c)+2be-(f+g)^2}} \Bigg) \Bigg/ \\
& 2\left(be^2+a(-bc+f^2)+g(-2ef+cg)\right)
\end{aligned}
\tag{17}
$$

Then the $j$-th component of the force on vertex $v_i$ is $f_{ij} = -\partial \hat{E}_{\text{excl}}/\partial x^j(v_i)$, and can be derived using the chain rule.

#### 5.2 Geometric interpretation of Eq. 17

Let $\iota$ be the denominator in Eq. 17, and then

$$
\begin{aligned}
\iota &= 2\left(be^2+a(-bc+f^2)+g(-2ef+cg)\right) \\
&= 2\left[\|r_{12}\|^2\langle r_{01}, r_{p0}\rangle^2 - \|r_{01}\|^2\left(\|r_{12}\|^2\|r_{p0}\|^2 - \langle r_{12}, r_{p0}\rangle^2\right)\right. \\
&\quad \left. -2\langle r_{01}, r_{12}\rangle\langle r_{01}, r_{p0}\rangle\langle r_{12}, r_{p0}\rangle + \|r_{p0}\|^2\langle r_{01}, r_{12}\rangle^2\right] \\
&= -2\langle (r_{01}\times r_{12}), r_{p0}\rangle^2
\end{aligned}
\tag{18}
$$

which is -72 times the square of volume $V$ of the tetrahedron formed by $x_0$, $x_1$, $x_2$ and $x_p$. Note that $\iota = 0$ when the four points lie in the same plane, but when the triangle is not degenerate, $I$ is only infinity when the point $x_p$ lies inside the triangle $t$.

Given the substitutions in Eq. 15, we can define some other values expressed using these substitutions.

$$
\begin{aligned}
h &= \|r_{02}\|^2 = a+b+2g \\
l &= \|r_{p1}\|^2 = a+c+2e \\
m &= \|r_{p2}\|^2 = a+b+c+2(e+f+g) \\
o &= \langle r_{12}, r_{p1}\rangle = f+g \\
p &= \langle r_{01}, r_{02}\rangle = a+g \\
q &= \langle r_{01}, r_{p1}\rangle = a+e \\
r &= \langle r_{02}, r_{p0}\rangle = e+f
\end{aligned}
\tag{19}
$$

The terms in the integral that show up in the square roots can also have geometric interpretation

$$
\begin{aligned}
\delta_2 &= ac-e^2 \\
&= \|r_{01}\|^2\|r_{p0}\|^2 - \langle r_{01}, r_{p0}\rangle^2 \\
&= \|r_{01}\times r_{p0}\|^2
\end{aligned}
\tag{20}
$$

$$
\begin{aligned}
\delta_1 &= ac + bc - e^2 - 2ef - f^2 + 2cg \\
&= c(a + b + 2g) - (e + f)^2 \\
&= ch - r^2 \\
&= \|r_{02} \times r_{p0}\|^2
\end{aligned}
\tag{21}
$$

$$
\begin{aligned}
\delta_0 &= b(a + c) + 2be - (f + g)^2 \\
&= bl - o^2 \\
&= \|r_{12} \times r_{p1}\|^2
\end{aligned}
\tag{22}
$$

And then the coefficients before each arctan can be further written as

$$
\begin{aligned}
\epsilon_2 &= -(af - eg)/\sqrt{\delta_2} \\
&= \|r_{01} \times r_{12}\| \frac{-\|r_{01}\|^2 \langle r_{12}, r_{p0}\rangle + \langle r_{01}, r_{p0}\rangle\langle r_{01}, r_{12}\rangle}{\|r_{01} \times r_{p0}\| \ \|r_{01} \times r_{12}\|} \\
&= 2A_t \cos\varphi_2
\end{aligned}
\tag{23}
$$

where $\varphi_i$ is the dihedral angle between $t$ and the triangle formed by $x_p$, $x_{i+1}$ and $x_{i+2}$ (where the plus in the subscripts takes the modulo of 3). Likewise,

$$
\begin{aligned}
\epsilon_1 &= (af + fg - be - eg)/\sqrt{\delta_1} \\
&= (-e(a + b + 2g) + (a + g)(e + f))/\sqrt{\delta_1} \\
&= (-eh + pr)/\sqrt{\delta_1} \\
&= 2A(t) \cos\varphi_1
\end{aligned}
\tag{24}
$$

$$
\begin{aligned}
\epsilon_0 &= (ab + be - g(f + g))/\sqrt{\delta_0} \\
&= (bq - go)/\sqrt{\delta_0} \\
&= 2A(t) \cos\varphi_0
\end{aligned}
\tag{25}
$$

Finally, the arctan terms can have their interpretation as well, using the fact that $\arctan(1/\tan(x)) = \frac{\pi}{2} - x$ when $x \in [0, \pi]$.

$$
\begin{aligned}
\zeta_2 &= \arctan\frac{e}{\sqrt{\delta_2}} - \arctan\frac{a + e}{\sqrt{\delta_2}} \\
&= \arctan\cot(\pi - \angle p01) - \arctan\cot\angle p10 \\
&= \angle p01 - \frac{\pi}{2} - \left(\frac{\pi}{2} - \angle p10\right) \\
&= -\angle 0p1
\end{aligned}
\tag{26}
$$

$$
\begin{aligned}
\zeta_1 &= \arctan\frac{e + f}{\sqrt{\delta_1}} - \arctan\frac{a + b + e + f + 2g}{\sqrt{\delta_1}} \\
&= \arctan\cot(\pi - \angle p02) - \arctan\cot\angle p20 \\
&= \angle p02 - \frac{\pi}{2} - \left(\frac{\pi}{2} - \angle p20\right) \\
&= -\angle 2p0
\end{aligned}
\tag{27}
$$

$$
\begin{aligned}
\zeta_0 &= \arctan\frac{f + g}{\sqrt{\delta_0}} - \arctan\frac{b + f + g}{\sqrt{\delta_0}} \\
&= \arctan\cot(\pi - \angle p12) - \arctan\cot\angle p21 \\
&= \angle p12 - \frac{\pi}{2} - \left(\frac{\pi}{2} - \angle p21\right) \\
&= -\angle 1p2
\end{aligned}
\tag{28}
$$

So Eq. 17 can be rewritten as

$$
\begin{aligned}
I &= \left(\sum_{i=0}^{2} \epsilon_i \zeta_i\right) / (-72V^2) \\
&= \frac{A(t)}{36V^2} \left(\cos\varphi_0 \cdot \angle 1p2 + \cos\varphi_1 \cdot \angle 2p0 + \cos\varphi_2 \cdot \angle 0p1\right)
\end{aligned}
\tag{29}
$$

And Eq. 12 would be

$$
\begin{aligned}
\hat{E}_{\mathrm{excl}} &= \frac{k_{\mathrm{excl}} A(t)^2}{18V^2} \left(\cos\varphi_0 \cdot \angle 1p2 + \cos\varphi_1 \cdot \angle 2p0 + \cos\varphi_2 \cdot \angle 0p1\right) \\
&= \frac{\pi k_{\mathrm{excl}}}{d_\perp^2} \eta(t, p)
\end{aligned}
\tag{30}
$$

where $d_\perp$ is the distance between point $p$ and the plane defined by triangle $t$, and $\eta(t,p) = (\cos\varphi_0 \cdot \angle 1p2 + \cos\varphi_1 \cdot \angle 2p0 + \cos\varphi_2 \cdot \angle 0p1)/(2\pi)$ can be considered as the "contribution factor" of the triangle inside the whole plane corresponding to the case $n = 4$. When the triangle grows infinitely large to span the whole plane, $\cos\varphi_i = 1$, and $\angle 1p2 + \angle 2p0 + \angle 0p1 = 2\pi$, so in this case the integral becomes $E_{\mathrm{excl}} = \pi k_{\mathrm{excl}}/d_\perp^2$, which is exactly Eq. 12 integrated over the whole plane.
